# Distribution and density of oxpeckers on giraffes in Hwange National Park, Zimbabwe

**DOI:** 10.1101/621151

**Authors:** Roxanne Gagnon, Cheryl T. Mabika, Christophe Bonenfant

## Abstract

Oxpeckers (*Buphagus sp*.) are two bird species closely associated to large mammals, including giraffes (*Giraffa camelopardalis*). We tested whether oxpeckers distributed themselves at random across individuals or aggregated on individual giraffes, and if birds select the host’s body parts with the expected greatest amount of ticks. By counting oxpeckers on giraffe’s body from photographs, we quantified the distribution of birds per hosts and over predefined zones on the giraffe body. Oxpeckers displayed a strong aggregation behaviour with few hosts carrying many birds while many carried a limited number or no birds, a pattern that was most exaggerated for males. Oxpeckers were disproportionately found on the mane and back, where the density of ticks is presumably the highest. This high aggregation level of birds is typical of parasitic species and could suggest that oxpecker distribution may mirror the distribution of ticks, their primary food resource, on giraffes. Abundance of ticks appears as a major driver of the oxpecker foraging behaviour, and the oxpecker–large herbivores system proves to be highly relevant for the study of host–parasite dynamics.

## 1 INTRODUCTION

The distribution of animals in the environment results from a complex sequence of behavioural decisions aiming at satisfying the energy requirements of individuals while minimizing costs of movements, competition with con-specifics or other species, in balance with the perceived risks of predation for prey species (Krebs, 1972). Variation in habitat suitability, in space and time, is of prime importance to the ecology of species with consequences on its distribution range (MacArthur, 1972), its mating tactic (Emlen & Oring, 1977), or its population dynamics (Lack, 1966; Ostfeld & Keesing, 2000). From an evolutionary perspective, animals should select habitats with the greater suitability (sensu Fretwell & Lucas, 1969), to maximize the individual fitness.

At the population level, when and where resources are available is a strong predictor of the abundance of animals. Assuming an homogeneous distribution of discrete resources patches in the landscape and a random walk of foraging animal trajectories, the expected number of foragers per patch is given by a Poisson distribution (Hutchinson & Waser, 2007). This simple model of encounters between motile entities underpins most multi-species interaction models, including the Lotka-Volterra (Lotka, 1956; Hutchinson & Waser, 2007) or Nicholson-Bailey models (May, 1978) for, respectively, predators–prey and hosts–parasitoid dynamics. Any deviation from the Poisson distribution is usually interpreted as a sign of aggregation (over-dispersion) or avoidance (under-dispersion) of individuals (Pielou (1969, p. 96) but see Taylor et al. (1979) or Sjö berg et al. (2000) for a discussion and alternatives). Several indices have been proposed to quantify aggregation levels among populations or species from count data (see Kretzschmar and Adler 1993 for a review), the most widely used being the aggregation index *k* (Shaw et al., 1998).

Aggregation may arise from the animal behaviour such as social interactions (Wittenberger, 1981), constrain on mobility among patches (Gueron & Levin, 1995), or if animals perform area-restricted search of food patches (Morales et al., 2010) or do copy what the other con-specifics do when using public information (Clark & Mangel, 1986).

In multi-species interactions such as in bird vs. large mammals associations, the distribution of the birds is first guided by the one of the mammalian hosts, conceptually equivalent to a resource patch. This scenario fits the association between oxpeckers *(Buphagus sp*.) and large herbivores, described in the early 20th century (Moreau, 1933). Yellow-billed (*B*. *africanus*) and red billed oxpecker (*B*. *erythrorynchus*) are two bird species of sub-Saharan Africa associated to savanna ecosystems (Hustler, 1987; Plantan, 2009; Palmer & Packer, 2018). Both species live and feed almost exclusively on the body of large herbivores such as African buffaloes (*Syncerus caffer caffer*), giraffes (*Giraffa camelopardalis*), black rhinoceros (*Diceros bicornis*), white rhinoceros (*Ceratotherium simum*), impalas (*Aepyceros melampus*), greater kudus (*Tragelaphus strepsiceros*), common elands (*Taurotragus oryx*) and sable antelopes (*Hippotragus niger*) (Hustler, 1987; Stutterheim et al., 1988; Palmer & Packer, 2018), but red-billed oxpeckers seem to favor smaller sized hosts. Oxpeckers mainly prey upon ectoparasites of their large mammalian hosts although they sometimes can snatch tissues from their host (Bezuidenhout & Stutterheim, 1980). In terms of resource selection, the foraging behavior of oxpeckers can be decomposed into two main steps. The first step for birds is to localize large mammals in the landscape which, for oxpeckers represent motile and widely dispersed resources patches of varying size. The second event takes place on the host’s body where oxpeckers will search for the most suitable body part in terms of ectoparasites. These sequential behavioural steps are organized according to hierarchical and complementary spatio-temporal scales (see Johnson, 1980), and drive the distribution of oxpeckers among host species (*e*.*g*. Diplock et al., 2018), among the different body part they forage on, down to the selected prey they feed on.

At a large spatial scale of observation, the distribution of oxpeckers among the different species of mammalian hosts has been documented for decades (Moreau, 1933; Grobler, 1980; Hustler, 1987; Ndlovu & Combrink, 2015) with a clear preferences for large-bodied species with high tolerance to the presence of birds (Diplock et al., 2018). The giraffe hence appears to be one of the key host for the two oxpecker species (Grobler, 1980; Veríssimo et al., 2017). Much less is known about how oxpeckers are distributed among individuals of a given host species, and reporting on maximum records *(e*.*g*. 51 birds on one side of a single giraffe, Veríssimo et al. (2017)) only gives a rough idea about this level of heterogeneity. Heterogeneity in the number of oxpeckers per hosts is however very likely as a recent comparison among large herbivores of different age and sex classes strongly suggests (Diplock et al., 2018). At the host level of resource selection (small spatial scale), oxpeckers have long been recognized to favor some body parts of their hosts (Mooring & Mundy (see 1996, for an example on impalas (*Aepiceros melampus*))) but what body parts is most attractive vary with the host species (Diplock et al., 2018; Palmer & Packer, 2018). For instance, Ndlovu & Combrink (2015) reported that red-billed oxpeckers were most frequently perch on the back and the head of buffaloes and white rhinoceros and that the neck was preferred on giraffes. Whether body part preferences of oxpeckers changes with individual host’s characteristics remains poorly known.

In this paper we investigated the among-host and within-host distribution of oxpeckers on giraffes at Hwange National Park, Zimbabwe. We extracted the number and location of yellow-billed (*Buphagus africanus*) and red-billed oxpeckers on *n* = 683 giraffes from 500 photographs collected since 2013. We aimed at testing the random encounter model between foragers and hosts, and the ideal free distribution (IFD, Fretwell & Lucas, 1969) with oxpeckers as foragers and giraffes as their primary large mammalian hosts. We tested the three following predictions:

1. *Distribution of oxpeckers among giraffes*: By comparing the observed distribution of the number of oxpeckers per giraffe with the expected prediction from theoretical models, one may infer the underlying behaviour of resource selection by birds and their movements. If oxpeckers search for giraffes at random in the landscape the random encounter model predicts a Poisson distribution of birds among hosts (Hutchinson & Waser, 2007). Alternatively, if oxpeckers aggregate preferentially on some particular giraffes because of marked differences in the parasite load or because of copying behaviour, the model predicts a negative binomial distribution of birds among hosts (Pielou, 1969);
2. *Distribution of oxpeckers on giraffe body*: According to the IFD (Fretwell & Lucas, 1969), oxpeckers should be distributed on the giraffe body parts proportionally to the local ectoparasite load. Therefore, if the ectoparasite density is homogeneous over the whole giraffe body, the IFD predicts a homogeneous number of oxpeckers per area unit. Alternatively, if ectoparasites concentrate on some specific body parts, the IFD predicts a heterogeneous distribution of birds, with higher densities of oxpeckers on giraffe body parts with the higher ectoparasite burden (Horak et al., 1983; Mysterud et al., 2014). From previous observations (Plantan, 2009; Ndlovu & Combrink, 2015), giraffe body parts with the highest oxpecker number should be, in decreasing order, the mane, the neck, the scapula and the back;
3. *Sex-differences in bird load of giraffes*: Many studies evidenced that the ectoparasite load is proportional to the body mass and skin surface of the host (Horak et al., 1983,1987; Koenig, 1997). Consequently a bigger host should carry more ectoparasites and hence, more birds than a smaller one. Sexual size dimorphism is observed among many species and is particularly observable between male and female giraffes, reaching a 43% difference for fully grown individuals (Shorrocks, 2016). We will therefore test the prediction that more birds are present on male than on female giraffes (Diplock et al., 2018);

## 2 MATERIAL AND METHODS

### 2.1 Study site

This study was undertaken around Main Camp in the northeast of Hwange National Park, the main biological reserve of Zimbabwe (HNP; 19°00’S, 26°30’E, extending from Main Camp to Giraffe’s Spring and Ngweshla pans; Fig. 1). This park covers 14 650 km2 and supports a population of approx. 2 800 giraffes (Shorrocks, 2016). The Bulawayo-Victoria Falls railway line defines the western boundary of HNP while border with Botswana draws the eastern boundary. The long-term mean annual rainfall is ca. 600 mm (CV = 25%) and generally falls between October and April to form seasonal wetlands. Because of this relatively low annual rainfall, a xerophile vegetation covers most of HNP. The woodland vegetation consists primarily of African teak (*Baikiaea plurijuga*) intersected with patches of camel thorn (*Acacia erioloba*) or leadwood (*Combretum imberbe*). Bushland savanna with patches of grassland makes 64% of the area, mainly around the many artificially maintained waterholes. HNP hosts many large and mega-herbivore species attractive to oxpeckers, including giraffes, plain zebras (*Equus quagga*), African buffaloes, wildbeest (*Connochaetes taurinus*), greater kudus, waterbuck (*Kobus ellipsiprymnus*) and impala, and the less abundant sable and roan antelopes (*H*. *equinus*).

**Fig. 1.**
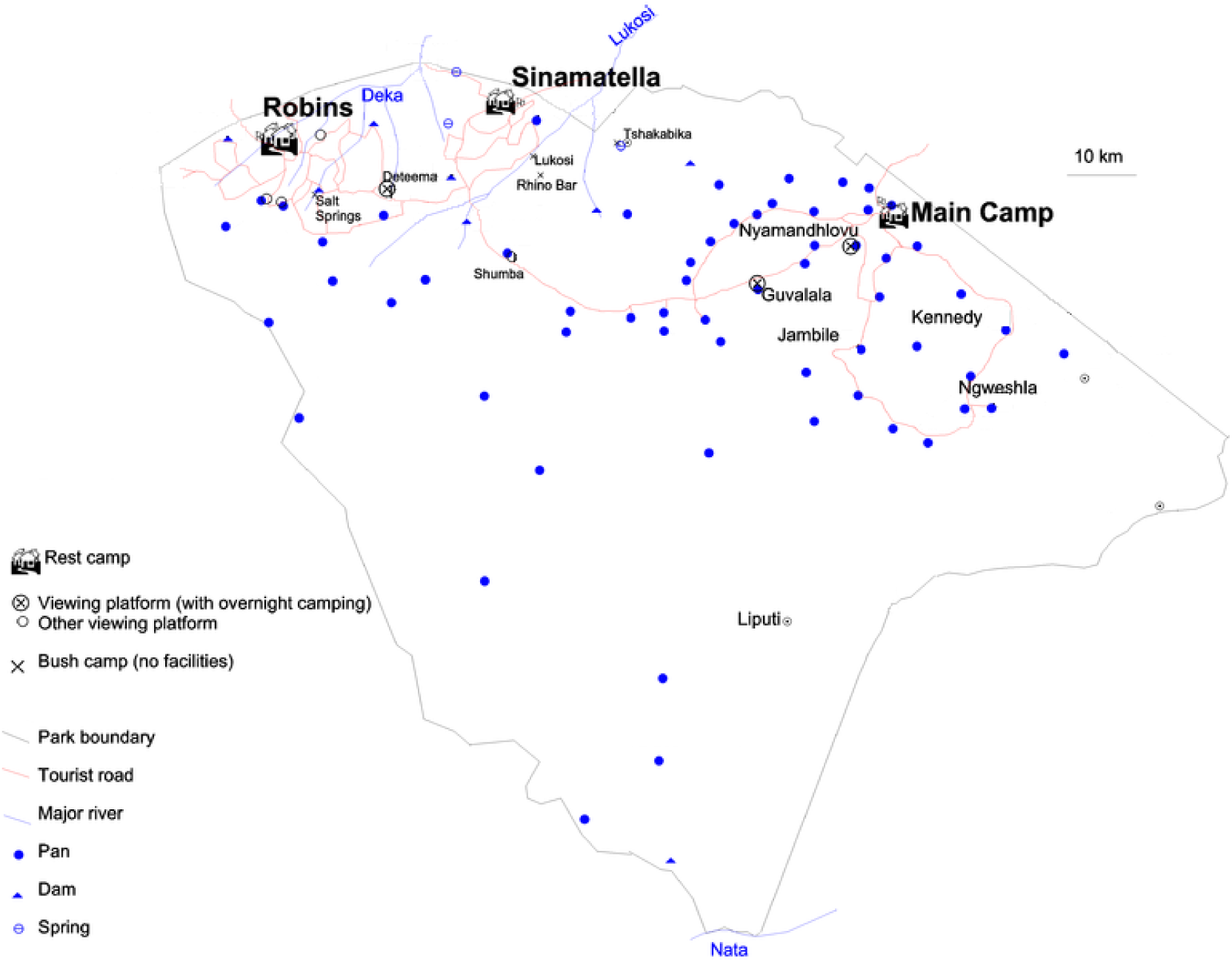
Hwange National Park map, Zimbabwe. The data collected (2013-2015) from oxpeckers (*Buphagus sp*.) and giraffes (*Giraffa camelopardalis*) derived from a study located in the Main Camp area, covering the north east of the park from Ngweshla to Giraffe Springs passing through Jambile. (‘Hwange National Park’ 2019, in Wikipedia: The Free Encyclopedia, Wikimedia Foundation Inc., viewed 11 May 2019, <https://en.wikipedia.org/wiki/Hwange_National_Park>)

### 2.2 Oxpecker biology

Red-billed oxpecker (RBO) and yellow-billed oxpecker (YBO) are two sympatric species, strictly african, which distribution ranges from Southern Africa up to Ethiopia on the East, with a few populations in Western Africa (del Hoyo et al., 2009). Although oxpeckers prey upon on ticks mainly, they can feed on wounded tissue, mucus, saliva, ear wax, hair and blood regularly (Bezuidenhout & Stutterheim, 1980; Weeks, 1999; Plantan, 2009). Some authors definitely think that the interaction between oxpekers and mammalian hosts is parasitism, while others support mutualism even though they admit oxpeckers can cause injuries to hosts (van Someren, 1951; Samish & Rehacek, 1999). In fact the relationship between oxpeckers and hosts could be context-dependent, where birds can be opportunists under particular biotic and abiotic conditions (Moreau, 1933; Nunn et al., 2011; Plantan, 2009). For instance, a mutualistic relationship may develop when the ectoparasite load is high on the host, but oxpeckers may become parasitic when hosts carry few ticks with too many birds, and leads to numerous open wounds.

### 2.3 Data collection

Oxpecker data derived from the study of giraffe ecology carried out at HNP. What giraffe sub-species (*sensu* Dagg, 2014) *currently live in HNP is not known but could be either G*. *c*. *angolensis* or *G*. *c*. *giraffa* according to the UICN (Muller et al., 2018). We have been monitoring the giraffe population opportunistically in 2012 and 2013, and on a regular basis since 2014, aiming at the photo-identification of individuals. Each year we drove the HNP road network daily (Fig. 1) for at least three weeks in a row and took photographs of every encountered giraffes with 200mm to 300mm lenses mounted on Nikon DSRL cameras. For all encounters, we recorded the date, the location and the time of observation along with group size and composition the individuals belonged to. We sexed giraffes based on the presence of bare skin at the ossicone tip for males, and on shape of the skull or visible male genitals when possible. We classified giraffes into four age-classes (calf, juvenile, sub-adult and adult) assessed in the field from their size and coat color since giraffes darken with age (Dagg, 2014). In this study we used photographs taken between 2013 and 2015, yielding a total of 500 photographs and, because several individuals were on the same frame sometimes, a sample size of 683 giraffes. Although we avoided to analyze a sequence of continuous photographs, re-observations of the same individual in time might have occurred, either during the same field session or from one year to another. For a given field session, if the same individuals are likely several days apart on average, we cannot exclude a few cases where the same giraffe has been seen on consecutive days (C.B pers. obs.). Doing so, we assumed little consequences of pseudo-replication on the results because of the rapid exchanges and movements of oxpeckers among hosts. We then counted oxpeckers on every giraffes seen on photographs and assigned each detected bird to one of the 14 predefined parts of the giraffe body (see Fig. 3A). Giraffe body parts were chosen so that they could easily be identified from landmark points whatever the point of view. From a flattened model of giraffe body (Fig. 3A), we approximated the relative area for the 14 considered parts with the *ImageJ* software. Relative surface ranged from < 1% for armpits and groins together, to 14% for the neck, for instance.

We excluded approx. 50% of all photographs because giraffes were too distant to reliably spot oxpeckers which roughly corresponded to a giraffe relative size smaller than ⅓ of the photograph height, or because of a poor image quality. Individuals whose whole body was not visible were rejected. For a subset of photographs we repeated oxpecker counts twice with two different observers (RG and CB) to estimate bird detection probability. Despite yellow-billed and red-billed oxpecker do occur at HNP, we did not differentiate betwen the two species in our analyses. The main reason for this choice was the difficulty to identify the species correctly from photographs when the bird’s beak was not clearly visible. It seems, however from the easiest cases, that the yellow -billed oxpecker is more abundant than the red-billed one (62 vs. 38% of our observation respectively).

### 2.4 Data analyses

We first estimated the detection probability of individual oxpeckers from photographs by setting a double-observer experiment (Nichols et al., 2000). The two observers reported the total number of detected birds they found (noted *x*_11_ and *x*_22_) from which we calculated the number of birds seen by observer 1 and missed by 2 (*x*_12_) and conversely (*x*_21_). The double-observer method returns the detection probability of observer 1 and 2 (respectively *p*_1_ and *p*_2_), as well as the average detection probability of birds *p*. The estimated number of oxpeckers per giraffe is then given by 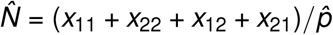. We fit the double-observer model to our data with the *unmarked* library (Fiske & Chandler, 2011) and tested for the effect of host sex on the detection probabilities.

We derived the aggregation index *k* from a particular parameterization of the negative binomial distribution (where the variance *V* is related to the mean by *V* = *λ* + *λ*^2^*/k*) that takes values close to zero with increasing levels of aggregation. To compare with previous studies, we also estimated the preference of oxpeckers for given hosts with a preference index (PI), calculated as the number of hosts counted divided by the number of oxpeckers counted (Grobler, 1980). A PI of 5 means one bird is seen every fifth counted hosts on average. We computed confidence limits of point estimates of PI with a non-parametric bootstrap (Manly, 2007).

We estimated the relative distribution of oxpeckers on giraffe body parts with a multinomial logistic regression. This particular type of GLM estimates the 14 probabilities (noted *π*_*i*_) of birds to be located on each body part of the giraffe holding the constrain that 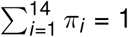. Because birds are more likely to be found on the largest body parts, we used the relative surface as an offset variable in the multinomial logistic regression to estimate the relative probability of use of each body parts. To control for the imbalanced number of males and female giraffes among years, we entered year as a categorical variable before testing for the effect the host sex (a 2-levels categorical variable) on the relative density of oxpeckers on body parts with likelihood ratio tests using the *nnet* library (Venables & Ripley, 2002).

Finally, we modeled the number of oxpeckers per giraffe with generalized linear mixed models (GLMMs). We tested for effects of host sex (a 2-levels categorical variable) and group size (continuous variable ranging between 1 to 5) while accounting for the potential confounding effects of year, season and daytime (hour extracted from the photograph metadata) on the number of seen birds with random effect variables. We fitted the GLMM and tested the effects of the explanatory variables with likelihood ratio tests using the *glmmTMB* library (Brooks et al., 2017). Our response variable being count data, we first fit a Poisson model but also considered the more flexible negative binomial distribution to account for potential over-dispersion of oxpecker counts (Ver Hoef & Boveng, 2007). We ran all analyses in the **R** 3.6.1 statistical software (R Core Team, 2018). Unless otherwise stated, we reported all estimated parameters as mean ± sd and predicted probabilities as the mean with its associated 95% confidence interval in brackets.

## 3 RESULTS

From a subsample of *n* = 117 giraffes, the overall detection probability of oxpeckers was 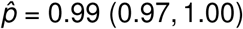 but differed substantially between the two observers (RG: 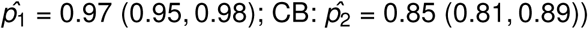. Although oxpeckers have 1.3 more chances to be seen on a male than on female giraffe, detection probability did not differ significantly with the host sex (estimated difference between male and female hosts on the logit scale: *β* = 0.26 ± 0.34, *P* = 0.45). On average we estimated oxpecker density to be *D*_*O*_ = 2.91 (2.63, 3.23) birds per giraffe once we accounted for imperfect detections. In the following analyses, RG did analyze all photographs.

When using *n* = 683 giraffes, mean oxpecker density was 2.16 ± 3.01 birds per host without accounting for detection probability. The overall preference index (PI) is 0.46 ± 0.10 with a maximum number of oxpeckers counted on a single host of 17 (Fig. 2). In support of the aggregation hypothesis, the estimated aggregation coefficient 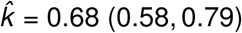 suggested a strong aggregation of oxpeckers on individual giraffes. Because the estimated aggregation coefficient *k* approaches zero, the negative binomial distribution converges to the logarithmic series distribution, with a strong skew toward giraffe carrying no bird (Fig. 2). Overall, our results lend support to the hypothesis of a non-random association between oxpeckers and giraffes at HNP (*H*_1_).

**Fig. 2.**
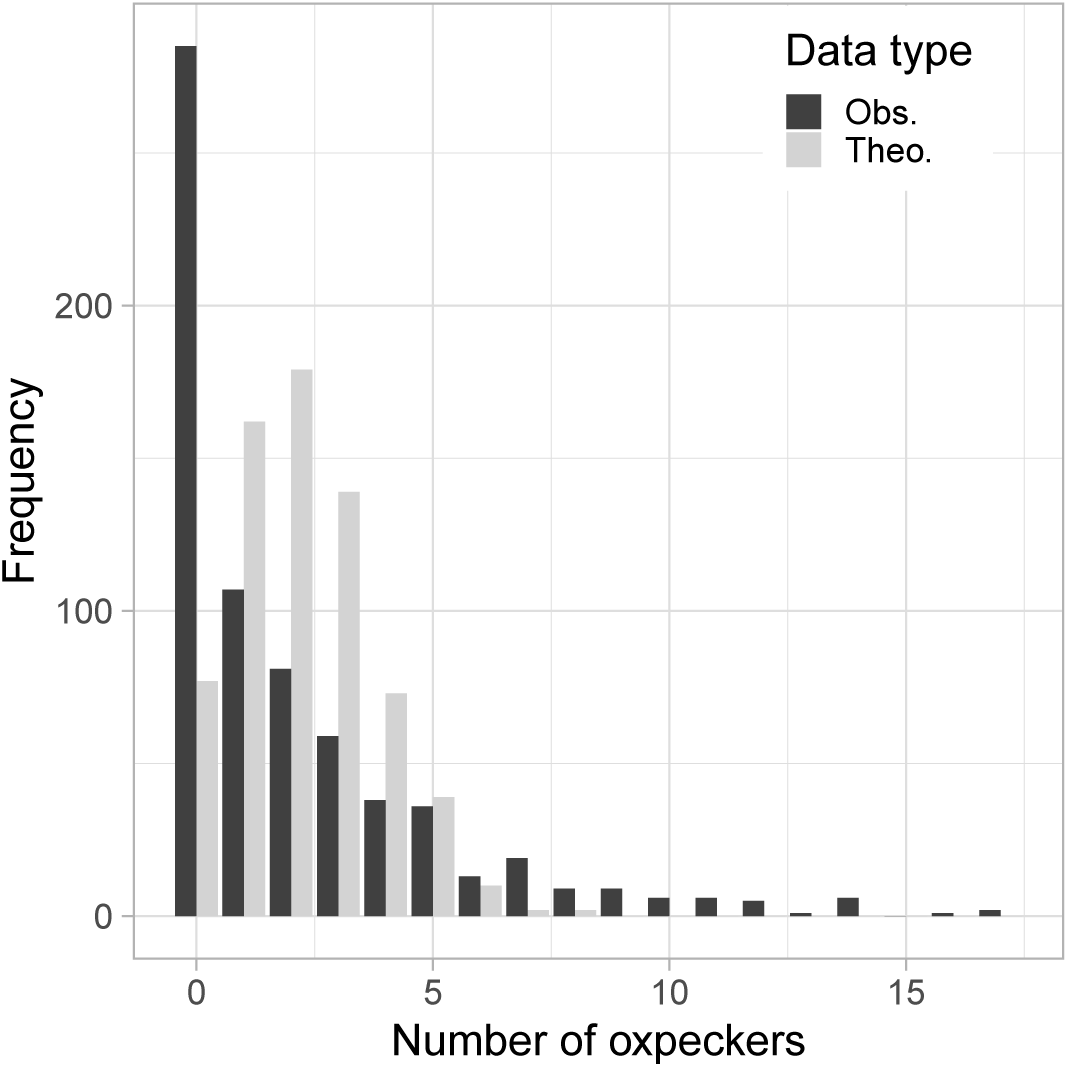
Distribution of oxpeckers (*Buphagus sp*.) on individual giraffes (*Giraffa camelopardalis*) at Hwange National Park, Zimbabwe. In black is the observed distribution and in grey the expected distribution according to a Poisson distribution law taking the observed mean as parameter (*λ* = 2.16). Note the marked over-representation of giraffes carrying no bird and the long distribution tail of giraffes with numerous birds on their body in the observed data.

**Fig. 3.**
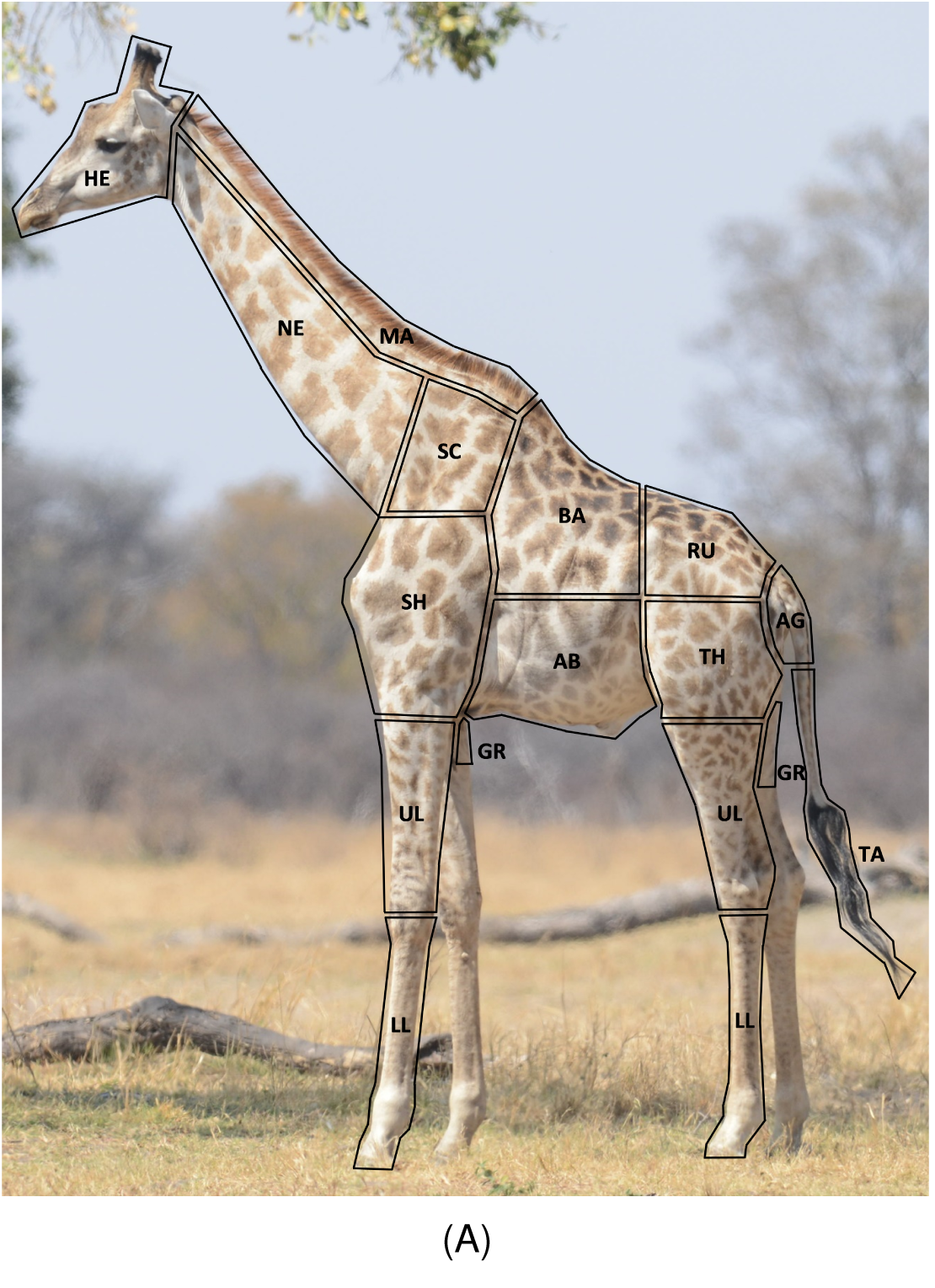
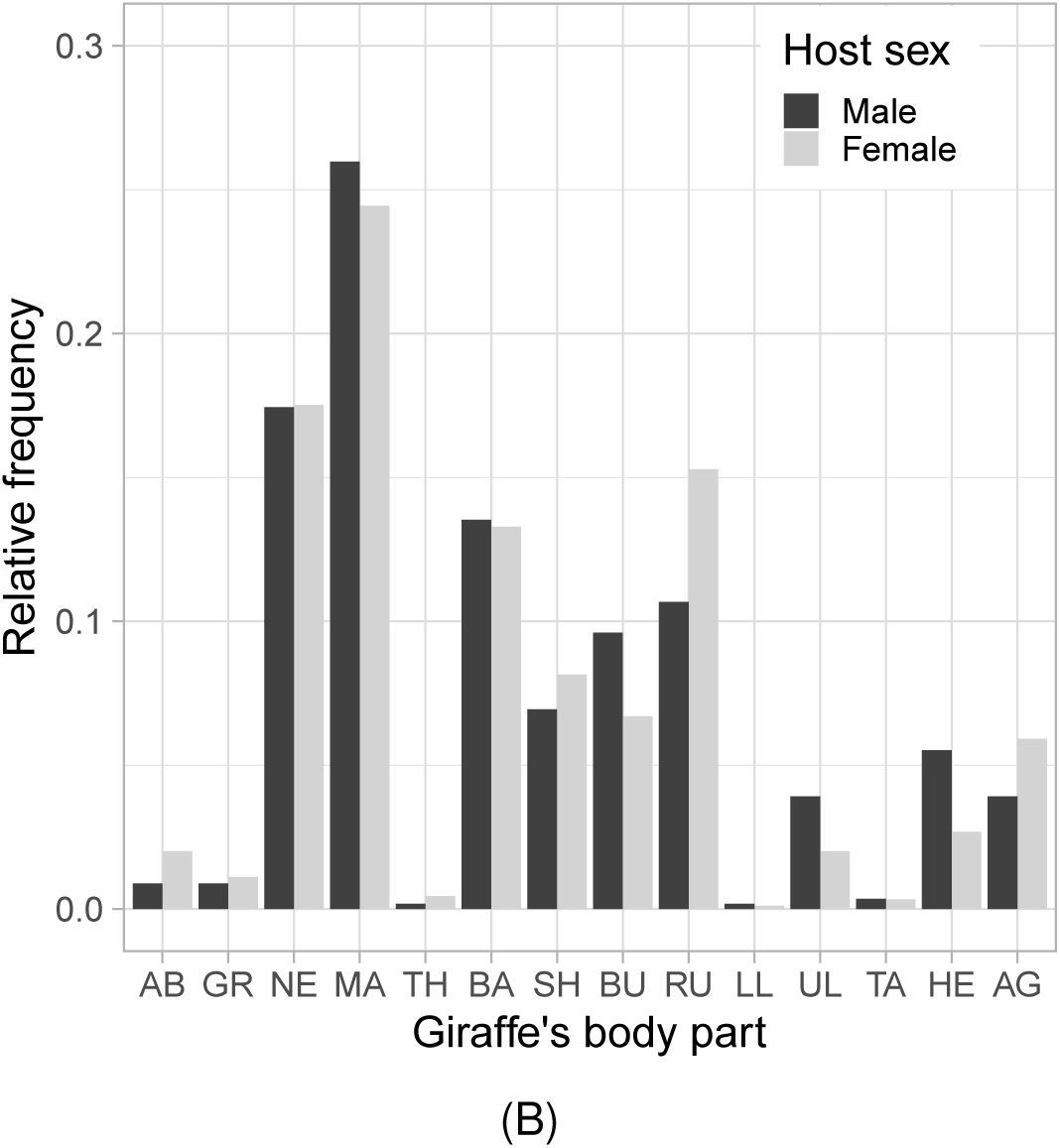
(A) Defined zonation and (B) proportion of oxpeckers (*Buphagus sp*.) counted on the 14 different body parts of giraffes (*Giraffa camelopardalis*) observed at Hwange National Park, Zimbabwe (*n* = 683 giraffes). The two-letter acronyms of the x-axis stand for: AB: abdomen, GR: groin, NE: neck, MA: mane, TH: thigh, BA: back, SH: shoulder, RU: rump, SC: scapula, LL: lower leg, UL: upper leg, TA: tail, HE: head, AG: ano-genital.

The relative distribution of oxpeckers on giraffe’s body deviated strongly from uniformity with some body parts being much more used than others (Fig. 3B). Supporting our hypothesis H_2_, birds gathered principally on the neck (*π* = 0.18 (0.15, 0.20)) and mane (*π* = 0.25 (0.21, 0.27)) of giraffes, but were rarely seen on the lower limbs (*π* < 0.01) or on the tail (*π* < 0.01). Oxpeckers did not use the giraffe’s body differently according to the host sex (likelihood ratio test: *χ*^2^ = 28.68, df = 13, *P* = 0.07; Fig. 3B), although relatively more birds used the ano-genital (*β* = 0.40 ± 0.56, *P* = 0.47) and the scapula areas (*β* = 0.45 ± 0.53, *P* = 0.39) of females compared to males. Conversely more birds were seen on the head (*β* = 1.53 ± 0.57, *P* < 0.001) and rump (*β* = 1.17 ± 0.54, *P* = 0.03) of male giraffes. Our results hence confirm the marked heterogeneous distribution of oxpeckers on the body of large mammalian hosts.

Although GLM with a Poisson distribution and a logarithmic link are usually recommended for count data (Agresti 2002), a preliminary goodness-of-fit (GOF) test suggested an over-dispersion of the data compared to a Poisson distribution (*χ*^2^ = 2877.21, df = 682, *P* < 0.001). A GLM a with negative binomial distribution did fit the data better than with a Poisson distribution (GOF test: *χ*^2^ = 609.10, df = 682, *P* = 0.98). The number of oxpecker per host decreased with group size for male giraffes (*β* = *-*0.23 ± 0.13, *χ*^2^ = 3.10, df = 1, *P* = 0.07; Fig. 5) but not for females (*β* = 0.07 ± 0.06; interaction sex by group size: *β* = *-*0.34 ± 0.13, *χ*^2^ = 6.53, df = 1, *P* = 0.01). As expected from our last hypothesis H_3_, the number of oxpeckers was larger on the giraffes exposing the largest body area to the birds (Fig 4). Accordingly, we found that oxpeckers were 20% more numerous on males than on females (density of 2.57 ± 0.23 and 1.97 ± 0.14 birds per giraffe respectively: *β* = 0.23 ± 0.12, *χ*^2^ = 3.75, df = 1, *P* = 0.05). The sex-specific aggregation coefficient reads *k* = 0.85 ± 0.09 and *k* = 0.62 ± 0.07 for females and males respectively, and was significantly smaller for male giraffes (bootstrap test: *β* = 0.23 ± 0.13, *P* = 0.02). Similar to the mean, relative variability in the number of oxpeckers per host was slightly larger for male (CV = 1.49) than for female giraffes (CV = 1.29) (see also Fig. 4).

**Fig. 4.**
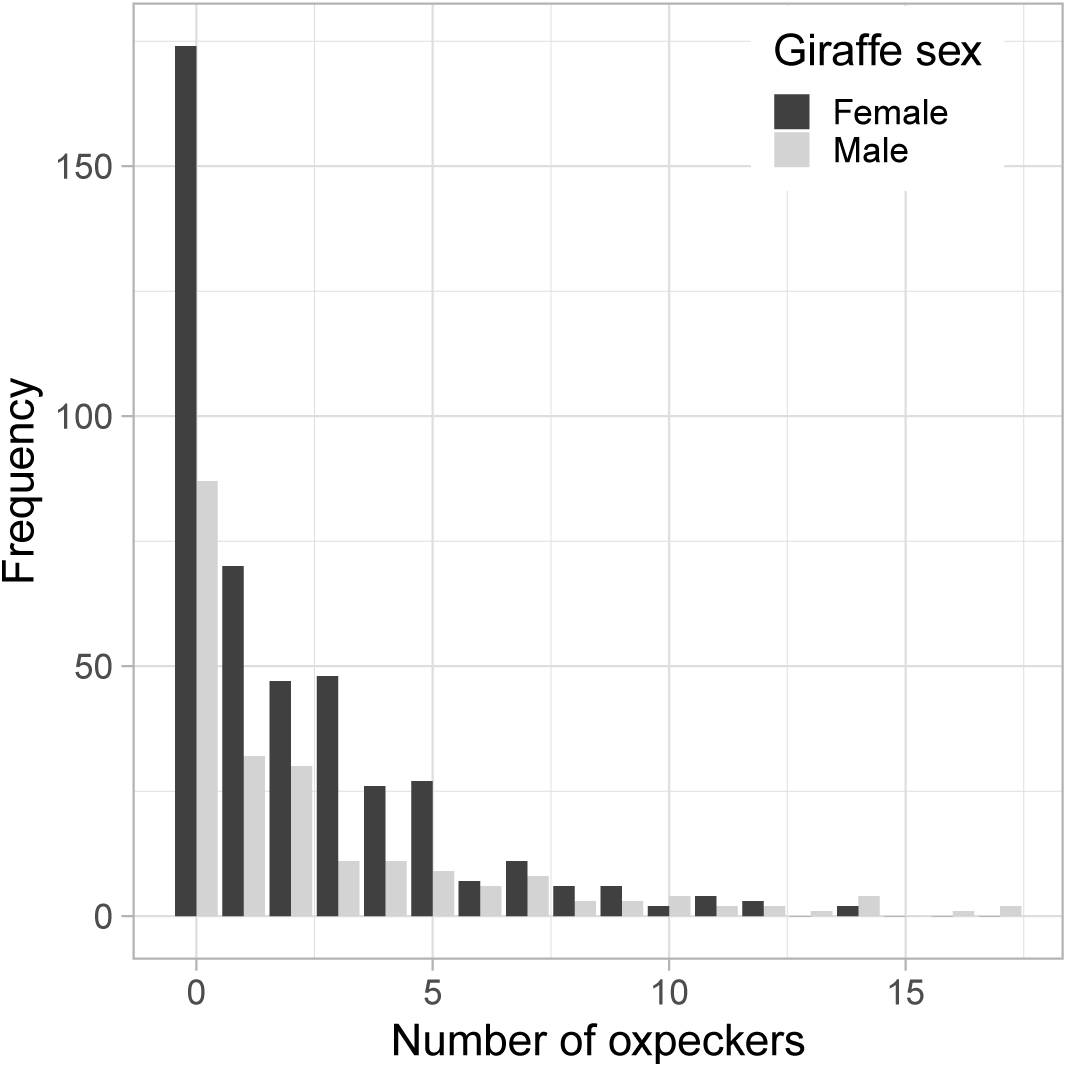
Distribution of oxpeckers (*Buphagus sp*.) on male and female giraffes (*Giraffa camelopardalis*) at Hwange National Park, Zimbabwe. Curves are the predicted frequencies as given by a negative binomial distribution model which parameters have been estimated separately for the two sexes. Vertical dashed lines represent the mean number of oxpeckers carried by individual giraffes. Note that males with no oxpecker are less frequent than females, and that the largest aggregations of oxpeckers have been found on male giraffes.

**Fig. 5.**
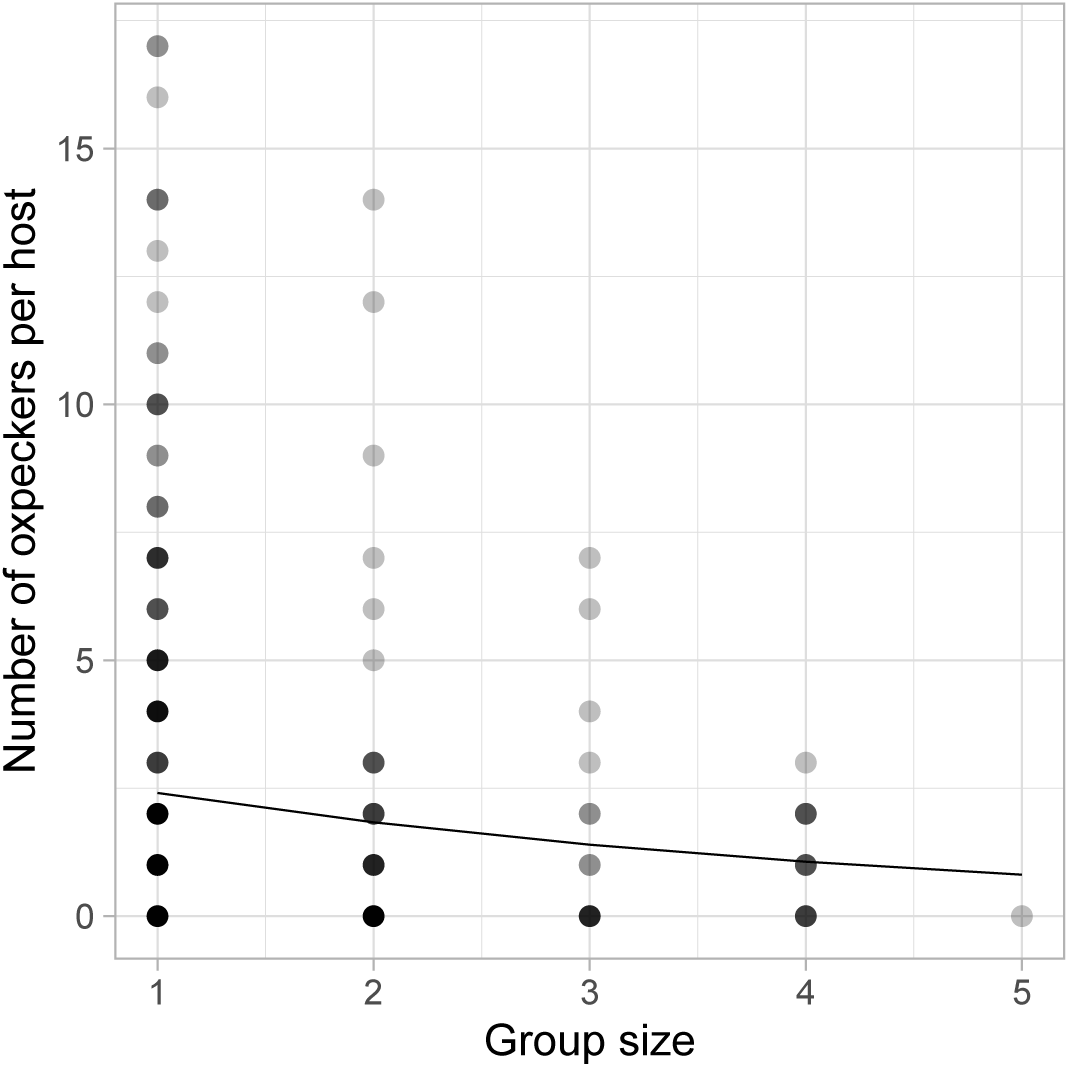
Variation in the number of oxpeckers (*Buphagus sp*.) per host according the group size of male giraffes (*Giraffa camelopardalis*) observed at Hwange National Park, Zimbabwe. Color density of dots is proportional to the number of observations (darker dots mean several records). The continuous black curve is the predicted density of oxpeckers by a generalized linear model with a negative binomial distribution (*β* = *XX* ± *YY, P* < 0.001).

## 4 DISCUSSION

The foraging behaviour and type of interaction between oxpeckers and their large mammalian hosts is poorly understood and still debated (Weeks, 2000; Nunn et al., 2011; Welsh et al., 2019). A closer look at the distribution of birds among and within giraffes at HNP clearly shows how heterogeneous it is at the host level with many carrying no bird while a few have *>* 10 birds on them. We also provide empirical evidences for non-random choice of host body part by oxpeckers, a behaviour likely driven by the amount of ticks birds can find and share with conspecifics. Overall the observed oxpecker distribution among giraffes at HNP matches with distributions generally observed in parasitic organisms (May, 1978), although we reckon it may only mirror the tick load of individual hosts.

At the largest spatio-temporal scale, oxpeckers have to chose among large herbivore species in the landscape, which is reflected by the host’s preference index (PI). The observed PI = 0.46 ± 0.10 for giraffes at HNP is similar to previously reported values in Kruger National Park (KNP), South Africa (0.90 for Grobler (1980); 0.54 for Ndlovu & Combrink (2015); 0.51 for Welsh et al. (2019) in Kenya). Surprisingly, Hustler (1987) found PI of 5.39 and 6.71 for giraffes in two separate areas of HNP. Host availability should indeed influence oxpeckers’ choice because the decrease in abundance of a key host such as the giraffe may force birds to switch to another less preferred but more numerous host with little fitness costs (Pyke, 1984; Hustler, 1987; Welsh et al., 2019). Here, host size plays a major role in host detection in flight and giraffes – like other large mammalian hosts – are easier to detect compared with smaller species (Grobler, 1980; Koenig, 1997). This is the main reason why the key host of oxpeckers alternates between buffaloes (Hustler, 1987), white rhinoceros (Ndlovu & Combrink, 2015) and giraffes (Grobler, 1980; Ndlovu & Combrink, 2015). The past high PI of giraffes at HNP suggests that birds must have exploited others hosts in the 80ies such as black rhinoceros, white rhinoceros, roans and sables (see Hustler, 1987, for details). This interpretation is supported by change in the composition of HNP’s community of large herbivores over the last decades, notably with the recent loss of the white rhinoceros (Valeix et al., 2008), which is consistent with the low PI values we report here for giraffes.

Focusing on the choice of individual giraffes by oxpeckers, we found a marked asymmetric distribution of birds (exponential distribution) whereby many carried no bird and a few ones were seen with up to 17 birds simultaneously (Fig. 2). This non-random distribution of oxpeckers among individual giraffes likely results from aggregation behaviour (Palmer & Packer, 2018). The aggregation coefficient *k* we found for the oxpecker distribution at HNP, close to 0, is typical of parasitic infections where only a few individuals are massively infested (Shaw et al., 1998). That oxpeckers similarly aggregate on a few giraffes would suggest they behave like parasites with their host in agreement with previous studies (Plantan, 2009; Nunn et al., 2011). Birds could use public information like conspecific density to chose a giraffe in a group (Doligez et al., 2004). Because oxpeckers mostly feed on ectoparasites, the marked aggregation of birds could indirectly mirror the distribution of ticks among giraffes. In mammals, infestation is indeed highly variable among hosts (*e*.*g*. Shaw et al., 1998; Brunner & Ostfeld, 2008). For instance, in roe deer *(Capreolus capreolus*), bank vole (*Clethrionomys glareolus*) and mountain hare (*Lepus timidus*), most host individuals bear few ticks and only a few individuals bear many (Horak et al., 1983; Talleklint & Jaenson, 1997; Mysterud et al., 2014). From this hypothesis, one could make indirect inference on tick burden of individual giraffes from the number of hosted oxpeckers, given the birds distribute themselves according to the ideal free distribution.

Within giraffes, we clearly found preferences for some body parts by oxpeckers. At HNP, oxpeckers were mostly found on the neck and the back of giraffes followed by the head, the abdomen, the lower limbs and the tail (see also Plantan, 2009; Ndlovu & Combrink, 2015, for similar results). The mane seems the most preferred giraffes’ body part of oxpeckers (Koenig, 1997). This row of hairs seems to be a favourable habitat for ectoparasites by providing shelter from predators (Ndlovu & Combrink, 2015) although oxpeckers use a scissoring behaviour to easily pick parasites from the hairs Koenig (1997). It has been noted that oxpeckers gather at the bigger host’s mane to hide from predators or when they are alarmed, which may contribute to increase their number in this area. This behavior may reflect an important driver of oxpecker group dynamics. However, contrary to other smaller herbivores (see Warwick et al., 2016, on cattle and sheep), we expected a relatively low density of ticks on the giraffe’s head because they rarely feed directly on the ground (Seeber et al., 2012). Our results support this assertion but oxpeckers could sometimes forage the head seeking for other food resources such as saliva, mucus, earwax (Ndlovu & Combrink, 2015) or wounds. This also could be the case for female genitalia where oxpeckers can also feed on mucus and secretion of their hosts (Weeks, 1999; Plantan, 2009). The abdomen, groins and armpits, thighs and tail present the lowest density of oxpeckers. Unlike mane, these areas are parts that can be easily groomed by giraffes, depleting tick quickly and making this area less preferred for oxpeckers (Koenig, 1997; Ndlovu & Combrink, 2015). Assuming that abundance or presence of ticks is highest in the mane running on the neck and back of giraffes, our results would concur with the distribution of oxpeckers as predicted by the optimal foraging theory (Pyke, 1984).

Density of oxpeckers differed substantially according to the sex of the giraffe, with male hosts carrying 20% more birds than females (Fig. 4). That male large herbivores are more frequently used as hosts than females by oxpeckers has been also reported in Tanzania for cattle and, to a lower degree, for wild species (Diplock et al., 2018). We add to the these previous observations that heterogeneity in the number of oxpeckers per hosts is larger for male than female giraffes. The simplest explanation is the one of male giraffes being larger in size than females, more birds can feed on a male holding a constant *per capita* food rate. The large variability in male body size in giraffe populations implied by their long body development (Shorrocks, 2016, p. 74) is in line with the wider distribution in oxpecker number in males than in females. Besides, the negative relationship we found between bird density and the observed group size of males (Fig. 5) suggests an alternative yet simple explanation for why males would carry more birds than females. Like in many sexually dimorphic species, male giraffes live in smaller social groups than females (Dagg, 2014). When a lone male is detected by an oxpecker flock, all birds will forage on this single individual, while they will likely dispatch themselves on all group member is several giraffe are found in proximity.

From a functional viewpoint, the higher abundance of oxpeckers on male than on female giraffes could, more likely, proceed from a higher load in ectoparasites and hence, larger food resources for the birds (Mooring & Mundy, 1996). The effect of host sex on the ectoparasite load are however equivocal in mammals (Ferrari et al., 2004; Kiffner et al., 2013). For instance, Horak et al. (1987) reported more ticks (*Amblyomma hebraeum*) on male kudus, which could make the female less attractive to oxpeckers. Conversely, tick load was similar whatever the sex and age of red deer (*Cervus elaphus*) in Scandinavia (Mysterud et al., 2014). Male giraffes could carry more ticks than females because of intra-sexual fights for reproduction. Neck fights, opponent chasing and female mounting result many injuries and open wounds all over male’s body (Nunn et al., 2011). Being opportunistic feeders, oxpeckers would benefit from the higher wound- and tick-feeding opportunities on male giraffes (Plantan, 2009). Moreover, sexually active males have, in general, a weaker immunity than females (Foo et al., 2017), which could indirectly contribute to their overall higher tick load compared to females. Consequently, if oxpeckers feed on giraffe individuals based upon their tick load, our results support a more variable tick infestation among males than females because of their contrasting body condition or immune-system efficiency (Fig. 4).

To evaluate the reliability of the oxpecker detection and location on giraffes from photographs, we carried out a double-observer experiment on a sub-sample of our images. We found the overall detection of the birds from photos to be very high (99%) but ideally all should be analyzed by 2 observers. Although not perfect, one person only (RG) scrutinized the 500 photographs henceforth keeping the detection and condition of observation similar for the complete data set. A major advantage of oxpecker counts from photographs is to make the counts and analyses repeatable and, contrary to what one might think, the use of one side of giraffes to locate and to count birds is an advantage because it avoids the issue of double counting. That said, we acknowledge that the major bias of our study was the location where the giraffe photographs were taken *i*.*e*. mainly around the many artificially maintained waterholes where large herbivores come to drink. When a large herbivore stand on the shore of waterholes, oxpeckers often use it as platform to rest, to sunbath, and to reach water to drink as well (Stutterheim, 1976). Consequently, the maximum number of oxpeckers per giraffe may be higher than elsewhere in HNP. To avoid this bias some studies tend to limit counting within 500 meters of water points (Grobler, 1980) but because HNP is densely covered with trees, observations of giraffes away from waterholes remained very difficult. Finally, we did not differentiate between yellow- and red-billed oxpeckers which behaviour in terms of host and body part selection differ slightly (Péron et al. 2019). Because the yellow-billed oxpecker was most abundant at HNP, our results are likely governed the behaviour of this *Buphagus* species rather than of the red-billed oxpecker.

## 5 CONCLUSION

Our study puts forward that the distribution and abundance of oxpeckers were surprisingly heterogeneous among and within giraffes. Some host body parts are clearly preferred for foraging by birds such as the neck and the mane because those areas could be suitable habitats for ticks. Gregarious hosts (female giraffes, buffaloes) travel and forage as a group thereby increasing local abundance and transmission of ticks (Koenig, 1997) to which oxpeckers could be excellent control agents on wild large herbivores and on domestic ones too (Ndlovu & Combrink, 2015). From an ecological point of view, the oxpecker–large herbivores system proves to be highly relevant and useful for the study of host-parasite dynamics.

## Acknowledgements

We would like to thank Simon Chamaillé-James and Marion Valeix for their critical reading of the manuscript. Thanks are extended to Martin Muzamba and Trust Dube for their invaluable help in data collection at Hwange, and to the park authorities of Zimbabwe for their support. Thanks are also extended to Fred Bercovitch and two anonymous reviewers for their interesting and constructive comments.

